# The Xanthophyll Cycle balances Photoprotection and Efficiency in the seawater alga Nannochloropsis oceanica

**DOI:** 10.1101/2024.10.31.621405

**Authors:** Tim Michelberger, Eleonora Mezzadrelli, Alessandra Bellan, Giorgio Perin, Tomas Morosinotto

## Abstract

Photosynthetic reactions require continuous modulation to respond to highly dynamic environmental conditions. Regulation of photosynthesis involves various mechanisms, which differ across phylogenetic groups. One such mechanism, found widespread in photosynthetic eukaryotes, is the xanthophyll cycle, which involves the reversible light-dependent conversion between the carotenoids violaxanthin, antheraxanthin, and zeaxanthin.

In this study, we investigated the impact of the xanthophyll cycle in *Nannochloropsis oceanica*, a seawater microalga member of Eustigmatophyta that features a peculiarly high content of xanthophylls. We generated and characterized lines with increased levels of the enzymes involved in the xanthophyll cycle, i.e. violaxanthin de-epoxidase (VDE) and zeaxanthin epoxidase (ZEP) and demonstrated that their content is a main factor in controlling the overall reaction rates and the dynamics of the xanthophyll cycle. Subsequent differences in the xanthophyll profile affect the activation of photoprotection mechanisms such as non-photochemical quenching and tolerance to reactive oxygen species. Interestingly, overexpression of VDE expands the limits of high light tolerance, whereas the increased content of ZEP facilitates faster recovery after exposure to light but also heightened photosensitivity under some conditions.

These findings underscore the critical role of the xanthophyll cycle in the regulation of photosynthesis in *Nannochloropsis,* where it is not simply a mechanism to respond to excess illumination, but plays a central role in modulating photosynthesis, fulfilling the complex task of balancing photoprotection and light-use efficiency under different environmental conditions.

## Introduction

Photosynthetic organisms are the main primary producers in most ecological niches, and their ability to exploit sunlight to drive the fixation of CO_2_ in biomass directly or indirectly supports most living organisms. Approximately half of global primary production is associated with aquatic environments and is highly dependent on microalgae, making these organisms essential for life (de Vargas et al., 2015).

In the natural environment, light energy absorbed by photosynthetic pigments, such as chlorophyll (Chl), can often exceed the metabolic capacity of the cell, driving the overreduction of the photosynthetic electron transport chain and consequently the generation of toxic reactive oxygen species (ROS). Photosynthetic organisms have evolved several mechanisms that regulate light-use efficiency and photosynthetic electron transport to lower the probability of overreduction and ROS damage (Li et al., 2009). Therefore, these mechanisms have a seminal influence on the dynamics of primary productivity in natural ecosystems, as well as on the light-conversion efficiency in crops (Ort et al., 2015). Photosynthetic organisms show diversity of regulatory mechanisms depending on their evolutionary history and adaptation to environmental conditions, and the exploration of this biodiversity is a valuable strategy to increase our knowledge of their biological relevance (Alboresi et al., 2019).

A major role for the modulation of photosynthesis is played by the xanthophyll cycle (XC), i.e. the light-induced mutual conversion of the di-epoxide xanthophyll violaxanthin (Vx) into the epoxide-free zeaxanthin (Zx) via the intermediate antheraxanthin (Ax), which is found in many different photosynthetic eukaryotes, such as plants, green algae, and diatoms (Coesel et al., 2008; Dautermann and Lohr, 2017). The xanthophyll cycle is regulated by the two enzymes <u>v</u>iolaxanthin <u>de-</u> epoxidase (VDE) and <u>z</u>eaxanthin <u>ep</u>oxidase (ZEP), both belonging to the lipocalin-like protein family characterized by a catalytic β-barrel domain (Frenette Charron et al., 2005).

Under light excess, the xanthophyll cycle is activated by a decrease of the pH in the thylakoid lumen caused by an imbalance between proton translocation associated with photosynthetic electron transport and its consumption by ATPase. The decrease in lumenal pH activates VDE, localized in the thylakoid lumen, which undergoes a conformational change (Arnoux et al., 2009) and associates with the thylakoid membrane. The substrate violaxanthin is found free in the thylakoid membrane (Hager and Holocher, 1994), after being released from antenna complexes (Morosinotto et al., 2002). Structural and *in vitro* studies suggest that VDE is active as homodimers (Arnoux et al., 2009; Saga et al., 2010; Fufezan et al., 2012) or oligomers (Hallin et al., 2016). For membrane association, VDE is dependent on the presence of nonbilayer-forming lipids such as MGDG (Sen and Hui, 1988; Latowski et al., 2004), the main lipid in the thylakoid membrane (Hölzl and Dörmann, 2019). MGDG is also necessary for the solubilization of xanthophylls in the lipid phase (Goss et al., 2005) and affects membrane fluidity, which in turn can control Vx de-epoxidation (Latowski et al., 2002). For the de-epoxidation reaction, VDE requires ascorbic acid (Asc) as an electron donor (Hager, 1969), whose availability was reported to correlate with VDE activity in plants (Müller-Moulé et al., 2002; Tóth et al., 2011).

When light is not in excess, VDE is deactivated and zeaxanthin is converted back to violaxanthin by ZEP, which requires NAD(P)H and O2, as an well as FAD as essential cofactor (Büch et al., 1995). Plants and green algae commonly rely on a single ZEP, while diatoms, which additionally contain a diadinoxanthin cycle, possess three ZEPs with partially complementary functions (Niyogi et al., 1997, 1998; Coesel et al., 2008; Dambek et al., 2012; Eilers et al., 2016; Liu et al., 2023). In *Arabidopsis*, ZEP is associated with the thylakoid membrane and is located in the stroma in equal portions (Schwarz et al., 2015; Bethmann et al., 2019). In plants, ZEP activity was proposed to be controlled through the chloroplast redox state and inhibited at high light, however, its exact regulation remains elusive (Naranjo et al., 2016; Da et al., 2018; Bethmann et al., 2019).

The accumulation of zeaxanthin contributes to photoprotection by directly scavenging Chl triplets and ROS produced during light excess, consequently reducing photodamage (Havaux and Niyogi, 1999). Zeaxanthin accumulation also enhances the activation of nonphotochemical quenching (NPQ), which quenches Chl excited states (i.e. Chl singlets), dissipating excess energy as heat, thus reducing the probability of generating ROS. In eukaryotes, NPQ also depends on the activity of specific molecular activators, namely PsbS and/or LHCSR/LHCX, depending on the species (Li et al., 2000; Peers et al., 2009; Alboresi et al., 2010; Bailleul et al., 2010).

NPQ and the xanthophyll cycle play an important role in protecting the photosynthetic apparatus from excess irradiation, and plants depleted of VDE showed increased susceptibility to high light and reduced fitness under natural conditions (Kulheim et al., 2002). Modulation of XC dynamics was also shown to be effective in improving productivity in plant crops (Kromdijk et al., 2016; De Souza et al., 2022) and in microalgae (Perin et al., 2023).

Although widely spread among photosynthetic eukaryotes, the impact of the xanthophyll cycle on photosynthesis regulation and NPQ is diversified among photosynthetic organisms (Niyogi et al., 1998; Pinnola et al., 2013; Quaas et al., 2015). To broaden our understanding of this diversity, here we investigated the impact of the xanthophyll cycle on photosynthesis regulation in the marine microalga *Nannochloropsis oceanica*, which belongs to the Eustigmatophyceae within the class of secondary endosymbiotic heterokonts. *Nannochloropsis* genus present only Chl *a*, lacking any other accessory Chl but especially have the unique feature of presenting violaxanthin as the most abundant carotenoid species (Basso et al., 2014; Litvín et al., 2016; Liu et al., 2023). Here we demonstrated that the levels of VDE and ZEP activity in *Nannochloropsis* control the dynamics of the xanthophyll cycle, and that the latter plays an important role in photosynthesis regulation, even more prominent than in other species. The decrease in zeaxanthin accumulation resulted in increased photosensitivity while overaccumulation affected light-use efficiency, showing that photosynthetic organisms face a trade-off between photoprotection and efficiency in the regulation of photosynthesis.

## MATERIALS AND METHODS

### Strains and cultivation conditions

*N. oceanica* tdTomato (Südfeld et al., 2022) was kindly provided by Sarah d’Adamo (Wageningen University, The Netherlands). In this study*, N. oceanica* ZEP1-OE, VDE-OE and Z1-V-OE were generated as described below. For liquid cultures, microalgae were grown in sterile F / 2, composed of filtered sea water (salinity 32 g l^-1^), 40 mM Tris pH 8.0 supplemented with 0.75 g l^-1^ NaNO3, 0.05 g l^-1^ NaH2PO4, 3.15 mg l^-1^ FeCl3, 4.16 mg l^-1^ Na2EDTA, 0.01 mg l^-1^ CuSO4, 0.022 mg l^-1^ ZnSO4, 0.01 mg l^-1^ CoCl2, 0.18 mg l^-1^ MnCl2, 0.006 mg l^-1^ NaMoO4, 0.005 mg l^-1^ vitamin B12, 0.1 mg l^-1^ vitamin B1 and 0.005 mg l-1 biotin. Cells were maintained in 500 ml Erlenmeyer flasks with constant shaking at 100 rpm and illuminated under a 12:12 light: dark photoperiod at 100 µmol photons m-2 s-1, provided by an array of white LEDs. Cultures for transformation were grown in a Multicultivator MC 1000-OD system (Photon Systems Instruments, Czech Republic) at 100 µmol photons m-2 s-1 in F/2 enriched with Guillard’s (F/2) Marine Water Enrichment (Sigma). Three days before transformation, the cell density was set at OD750 = 0.1 and the culture was supplemented with 10 mM NaHCO3.

Solid medium was made from F/2 supplemented with Guillard’s (F/2) Marine Water Enrichment and 1% plant agar (Panreac AppliChem). For spot growth tests, algae were supplemented with 10 mM NaHCO3 and illuminated with the light protocols indicated in the respective figures under white LED light. For all treatments with high and fluctuating light, illumination was provided by a SL3500 LED light source (Photon Systems Instruments, Brno, Czech Republic). Spot tests were imaged with a Konika Minolta Bizhub C280 scanner and spot density was quantified with Fiji/ ImageJ. To test sensitivity to chemically induced oxidative stress, algae lines were inoculated in 24-well plates starting from the same OD750 (= 0.08) and cultured under constant illumination (100 µmol photons s^-1^ m^-2^) and in 1% CO_2_ (v/v). The formation of different ROS species was induced by adding methyl viologen or H2O2 at the indicated concentrations at the beginning of each experiment. Sensitivity to ROS was assessed by monitoring growth for 9 days by measuring OD750 with a Tecan Spark plate reader. For all growth conditions described above, the algae were grown at 22 ° C. Light intensities were defined using a LI-250A luminometer (Heinz-Waltz, Effeltrich, Germany) and cell numbers were determined with an automatic cell counter (Cellometer Auto X4 Cell Counter, Nexcelom).

### Generation of ZEP1-OE, VDE-OE and Z1-V-OE lines of Nannochloropsis oceanica

ZEP1-OE, VDE-OE and Z1-V-OE lines were generated by homologous recombination in a *N. oceanica* tdTomato landing pad strain as previously described by Südfeld et al. (2022). Sequences encoding ZEP1 (Gene ID: KU980906.1/ NO07G03040.1) and VDE (Gene ID: KU980905.1/ NO24G00840.1) were amplified from *N. oceanica* tdTomato cDNA and fused to the chloroplast target peptide of *N. oceanica* VCP1 (Gene ID: NO25G00860, Table S1). cDNA was obtained as described by Perin et al. (2023). The generated constructs were inserted into a transformation cassette that included a resistance cassette to Blasticidin S and homologous flanks for localization to the landing pad. 2.5 µg DNA were then transformed into *N. oceanica* tdTomato as previously detailed by Perin et al. (2015). Transformed cells were plated on F/2 supplemented with 100 µg/ml Blasticidin S and resistant colonies were isolated after 3-4 weeks.

### On-plate fluorescence screening

The transformants of *N. oceanica* tdTomato were tested for loss of tdTomato fluorescence directly on the plate using a stereomicroscope (Leica MZ16F). tdTomato fluorescence was imaged with an N2.1 LP filter and chlorophyll fluorescence was visualized with a chlorophyll LP filter. Images were captured with a 5-megapixel camera and analyzed with ImageJ to select colonies with tdTomato fluorescence loss.

### Genotyping of overexpression Strains

The successful transformation of *N. oceanica* tdTomato was verified by genotyping PCR. Genomic DNA was extracted by mixing a small algal sample with 5% Chelex-100 resin (BioRad), incubated at 95°C for 20 min and 4°C for 20 min. After vortexing for one minute and rapid centrifugation, the clear supernatant was used directly as the PCR template. The transformants were screened for the presence of the transgene and the integrated cassette using the primers listed in Table S2.

### Western blot

Western blot was used to detect proteins and their respective sizes. 5 x 108 cells were resuspended in 100 µl of B1 buffer (20 mM Tris-KOH pH 7.8, 400 mM NaCl, 2 mM MgCl2) for total proteome extraction or 75 mM Tris pH 8 for soluble proteome extraction, each supplemented with 3 mM benzamidine and 1 mM PMSF. The suspension was mixed with acid-washed glass beads (Sigma, G1145) and bead beaten 5 times for 20 s at 3500 rpm using a Bullet Blender Storm Pro homogenizer (Next Advance), with pauses of 3 min on ice. For total proteome extraction, cell lysates were solubilized in sample buffer (45 mM Tris pH 6.8, 30 mM DTT, 3% SDS and 10% glycerol), the beads were beaten again, incubated at room temperature for 20 min, centrifuged and the supernatant was transferred to a new tube. For the extraction of the soluble proteome, the lysates were incubated at -20°C for 30 min, centrifuged, and the supernatant was mixed with the sample buffer. Proteins in the extracts were then separated by SDS-PAGE, using homemade stacking gels (125 mM Tris pH 6.8, 4% acrylamide, 0.1% SDS, 0.6% TEMED and 0.1% APS) and running gels (1.24 M Tris pH 7.8, 12% acrylamide, 0.33% SDS, 0.7% TEMED and 0.1% APS). The gels were run at 50 V for 2.5 h in 250 mM Tris pH 8.3, 1.92 M glycine and 1% SDS. The separated proteins were transferred to a nitrocellulose membrane (Cytiva Amersham) in 20 mM Tris pH 8.3, 20% methanol and 152 mM glycine at 100 V for 1 h at 4°C. The membrane was blocked with 10% milk in TBS. Proteins with a 6xHis tag were detected with an α-His antibody (Novagen, #70796) in a 1:1000 dilution. VDE was detected with a homemade α-*A. thaliana-VDE* produced in rabbit and used in 1:500 dilution (Saga et al., 2010). Secondary HRP-conjugated antibodies against mouse (BioRad #1721011) and rabbit (Agrisera #AS09-60s) were used, and proteins were detected with Western HRP substrate (Immobilon Millipore, USA) and visualized with a CHEMI premium imager (VWR, Italy). The detected protein bands were quantified with the gel analysis tool in Fiji/ ImageJ.

### Pigment extraction

To quantify chlorophyll *a* and total carotenoid content, algal biomass was mixed 1:1 with N,N-dimethylformamide and incubated for 24 h at 4 ° C in darkness. The pigment contents were then quantified spectroscopically using specific extinction coefficients (Wellburn, 1994). To quantify individual xanthophylls after specific light treatments, 108 cells in 1 ml of supplemented F/2 were subjected to the desired light protocol and flash frozen in liquid nitrogen to capture a snapshot of the xanthophyll pool. For pigment extraction, cells were thawed on ice, washed once with deionized water, and resuspended in 50 µl 90% acetone. Acid-washed glass beads were added, and the sample was bead-beaten 4 times for 20 s at 3500 rpm, with pauses of 2 min on ice. 150 µl 90% acetone was added and another 5 s bead beating step was performed. The suspension was then centrifuged for 10 min at 10000 rpm, the supernatant was transferred to a fresh tube, and the extraction was repeated two more times. The extracted pigments were then analyzed by high pressure liquid chromatography (HPLC) as previously detailed by Perin et al. (2023).

### Chlorophyll fluorescence measurements

Photosynthetic metrics were evaluated from *in vivo* chlorophyll fluorescence measurements. On-plate chlorophyll fluorescence was assessed with a FluorCam 800 MF (Photons Systems Instruments, Brno, Czech Republic) as described by Perin et al. (2015). Liquid cultures were measured with a Dual PAM-100 fluorimeter (Heinz-Walz, Germany) as detailed in Perin et al. (2023). Before measurements, cells were dark-adapted for 30 min and exposed to the light protocols detailed in the respective figures. NPQ, the maximum quantum yield of PS II (ΦPSII) and the quantum yield of PSII (Φ’PSII) were determined as presented by Maxwell and Johnson (2000).

## RESULTS

### Generation of N. oceanica strains over-accumulating xanthophyll cycle enzymes

To assess the impact of an altered xanthophyll cycle, *Nannochloropsis oceanica* IMET1 lines overexpressing ZEP, VDE, and both simultaneously were generated using a recently developed strain exploiting RNA polymerase I to drive strong protein overexpression (Südfeld et al., 2022). Two ZEP-encoding genes are present in the genome of *N. oceanica* (Vieler et al., 2012). ZEP1 is closely related to ZEPs of diatoms, most green algae, and plants (Liu et al., 2023). Instead, ZEP2 forms a separate clade with ZEPs found in some green microalgae (Liu et al., 2023). Both enzymes were suggested to have complementary functions, ZEP1 being most expressed (Liu et al., 2023), while ZEP2 was found to be up-regulated under light stress (Wang and Jia, 2020). Here we focused on ZEP1 which was also recently demonstrated to play a more prominent role *in vivo* (Liu et al., 2023). The genes encoding VDE and ZEP1 (both tagged with the C-terminal 6xHis and c-myc Tag) were targeted for insertion by homologous recombination at a locus expressing a red fluorescent protein (tdTomato, Figure S1A, for simplicity the parental strain is hereafter named *N.oc.* WT). This enabled the identification of positive lines after transformation based on the loss of tdTomato fluorescence using a simple screening (representatively shown in Figure S1A-B). For the simultaneous overexpression of ZEP1 and VDE, the two overexpression cassettes were serially connected, and transgenic lines were generated as described above (Figure 1A).

**Figure 1.**
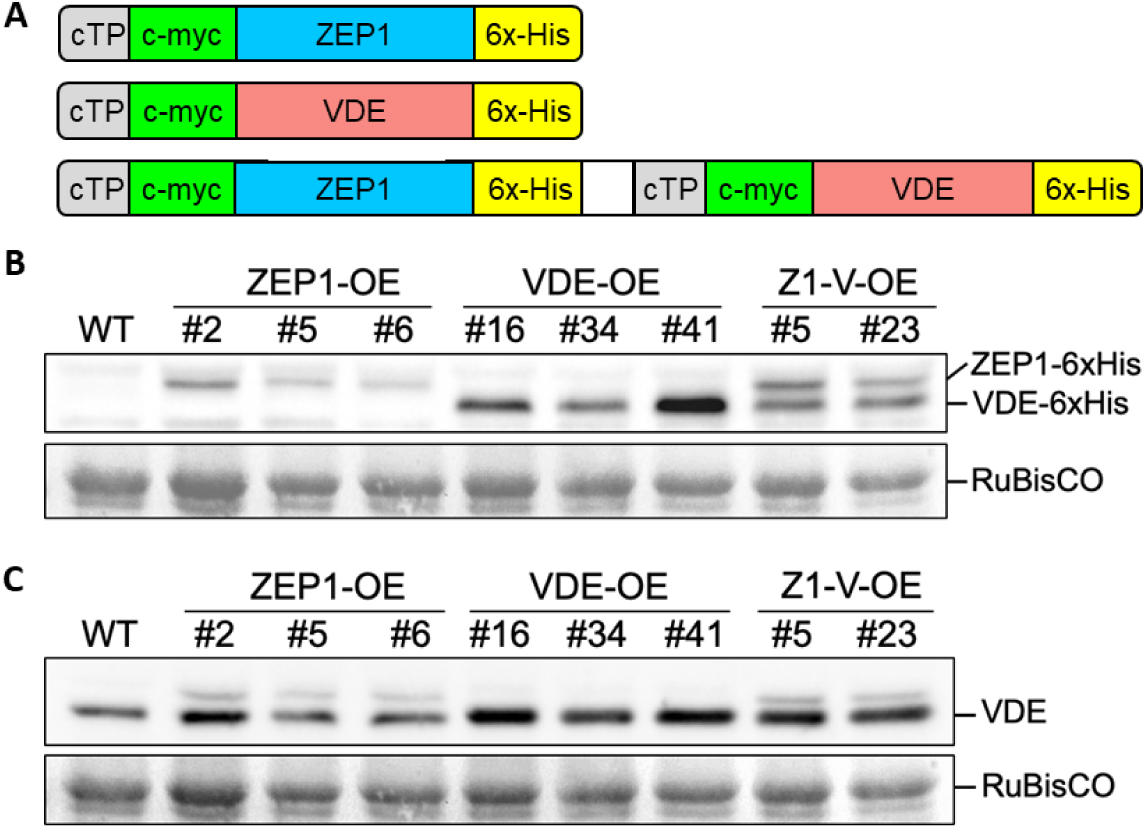
Isolation of *N. oceanica* strains overexpressing ZEP1/ VDE. **A)** Cartoon representation of constructs to generate overexpression of ZEP1 (upper panel), VDE (middle panel) and simultaneous ZEP1 and VDE overexpression (lower panel). eTP, chloroplast target peptide of VCP1; ZEP1, zeaxanthin epoxidase 1; VDE, violaxanthin de-epoxidase. B) Western blot showing ZEP1 / VDE overexpression in ZEP1-OE, VDE-OE and Z1-V-OE strains. Total proteome was extracted, separated in SDS-PAGE and analyzed with Western blot, using α-His antibody. Sizes of overexpressed constructs were determined with ProtParam from Expasy: ZEP1-6xHis, 54 kDa; VDE-6xHis, 49 kDa. C) Western blot showing VDE protein levels in N. *σc*. WT and the clones of the three newly generated lines. Total proteomes were separated in SDS-PAGE and analyzed with Western blot using α-VDE antibody. The α-VDE antibody also slightly detects ZEP1 resulting in a higher second band in ZEP1-0E and Z1-V-OE lines. RuBisCo levels are shown as loading control.

As overexpression of ZEP1 or VDE presumably affected the xanthophyll cycle and NPQ kinetics, lines showing loss of tdTomato fluorescence were further screened by NPQ analyses of isolated clones (see examples in Figure S1C-E). From ZEP1 or VDE overexpressors (ZEP1-OE / VDE-OE), more than 80% of the lines showing loss of tdTomato fluorescence also had altered NPQ activation. Three independent lines were randomly selected from those for further analysis.

For lines that simultaneously overexpressed both enzymes (Z1-V-OE), most of the lines selected by fluorescence screen showed a phenotype similar to VDE- or ZEP1-OE (Figure S1E). Instead, two lines showed an intermediate phenotype and were selected for further investigation.

In all selected independent lines, the insertion of transformed cassettes was validated by PCR (Figure S2). The accumulation of overexpressed ZEP1-6xHis and VDE-6xHis was confirmed using antibody against the 6xHis-tag, which showed protein accumulation in all lines, also revealing that the overexpression levels of each enzyme were similar in all individual and double overexpressors (Figure 1B, S3A-B). Interestingly, VDE was easily detectable after extraction of only soluble proteins, while ZEP1 was absent in this fraction and was detected only in total proteome extracts, suggesting that ZEP1 is associated with the thylakoid membrane in this species (Figure S4, 1B). Blotting with antibody against VDE from *A. thaliana* confirmed the previous results, also revealing that VDE-overexpressing cells accumulated ∼2-3x more VDE than WT cells (Figure 1C, S3C). The ZEP antibody raised on the *A. thaliana* protein did not recognize the algal isoform, while the α-VDE antibody also detected a weak band in ZEP1 overexpressing lines at the molecular weight expected for ZEP1 (Figure 1C). This weak recognition of ZEP1 can be explained by the fact that the α-VDE antibody was raised against the lipocalin domain of VDE (Saga et al., 2010), and VDE and ZEP1 are both members of the lipocalin protein family (Hieber et al., 2000) and share some sequence similarity.

### Impact of ZEP1/ VDE overexpression on xanthophyll cycle dynamics

We first investigated whether ZEP1/ VDE overexpression affected the overall composition of the photosynthetic apparatus by assessing the total chlorophyll and carotenoid content. In all generated lines, there were no relevant differences in the pigment composition except for slightly reduced chlorophyll levels in lines overexpressing VDE and a small increase in the Chl/ Car ratio in ZEP1-OE strains (Figure S5A-B).

The individual carotenoid fractions were dissected in the randomly selected clones of each overexpression lines using HPLC. *N. oceanica* cells showed detectable amounts of Zx and Ax, even when grown in dim light conditions, which is consistent with previous findings (Basso et al., 2014), and suggests that XC was already partially active (Figure 2A). Interestingly, in ZEP1-OE cells the basal amount of Zx was reduced, while it was increased in VDE-OE cells (Figure 2A), implying that overexpressed ZEP1 and VDE are active and can change the balance of XC towards epoxidated or de-epoxidated species, respectively, even during cultivation under low light. In Z1-V-OE cells, the composition of xanthophylls was indistinguishable from WT cells (Figure 2A), probably because simultaneous overexpression resulted in an equilibrium of the XC similar to the WT.

**Figure 2.**
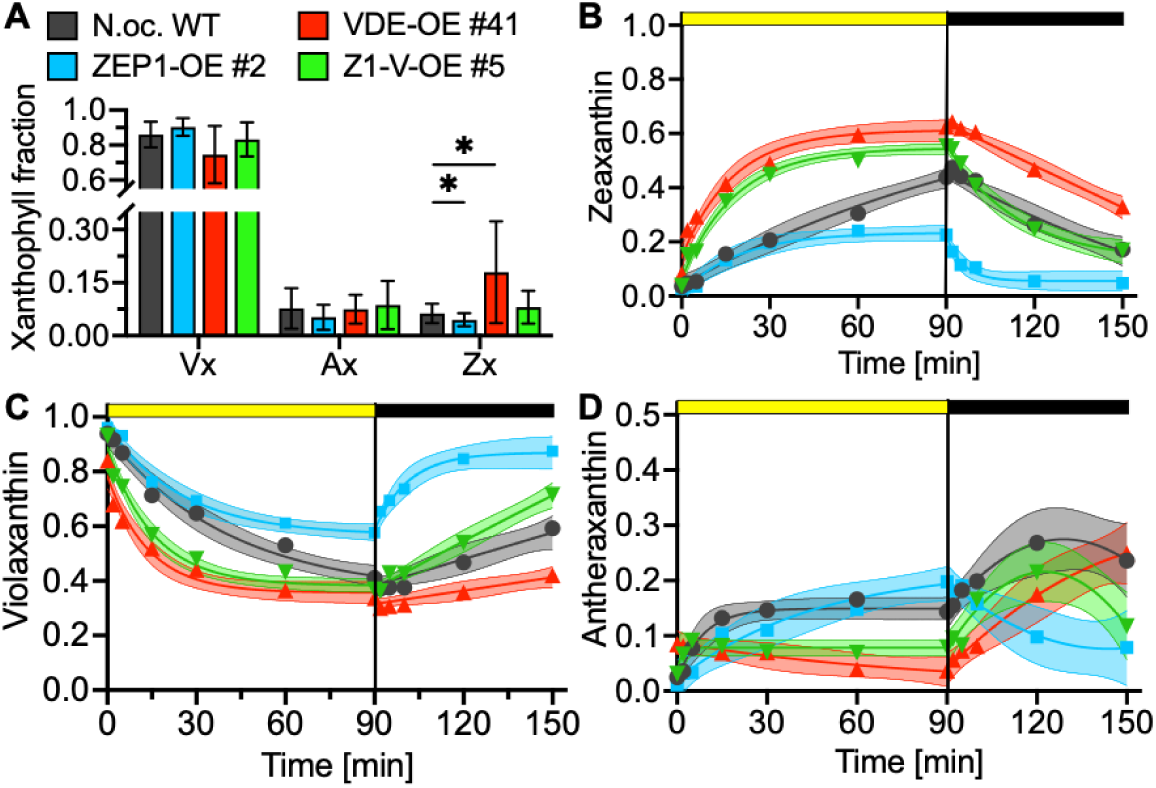
Influence of ZEP1/ VDE overexpression on xanthophyll cycle kinetics in *N. oceanica*. **A)** Xanthophyll fractins under standard growth conditions in a photoperiod of 12 h light (100 µmol photons s^-1^ m^2^) and 12 h darkness. Data represents means ± SD from ≥ 7 biological replicates. Significance was assessed with one-way Welch-ANOVA tests for each individual xanthophyll, *p < 0.05. **B-D)** Time-resolved analysis of xanthophyll fractions in light and darkness. Prior to measurements, cells were dark-adapted for 1.5 h to fully relax the XC. To follow XC activation cells were subjected to saturating light (1000 µmol photons s^-1^ m^2^) for 90 min. To follow XC relaxation, cells were subsequently exposed to darkness for 60 min. B) Zeaxanthin fractions. Data in light phase was fitted to an exponential plateau model. Data in dark phase was fitted to a one-phase decay model. C) Violaxanthin fractions. Data in light phase was fitted to a one-phase decay model. Data in dark phase was fitted to a second-order polynomial model. D) Antheraxanthin fractions. Data in light phase was fitted to an exponential plateau model. Data in dark phase was fitted to a second-order polynomial model. Data points in B-D represent means and curves are depicted as modelled fittings ± 95% confidence intervals from ≥ 3 independent biological replicates.

All OE lines showed no variation in vaucheriaxanthin esters (Vauchx) (Figure S6A-C), different from what Liu et al. (2023) reported for ZEP1-OE lines. For β-carotene we observed only minor alterations (Figure S6A-C), suggesting that ZEP1 and VDE overexpression caused only small, if any, differences in the composition of pigments not related to the XC.

Next, we sought to analyze the impact of ZEP1/ VDE overexpression on XC dynamics. For this, dark-adapted cells were exposed to saturating illumination (1000 µmol photons s-1 m-2) to induce XC activation and subsequently switched to darkness to follow XC relaxation. The dark-adapted cells overall showed reduced levels of de-epoxidated xanthophylls compared to cells grown in the light, because of the inactivity of VDE in darkness. Dark-adapted ZEP1-OE cells exhibited lower amounts of Ax and Zx, while VDE-OE cells showed higher levels of both compared to WT. Z1-V-OE cells again had a similar xanthophyll composition to the WT (Figure S6D).

Upon exposure to light, WT cells activated Zx synthesis at a relatively constant rate for 90 min. Similarly, ZEP1-OE cells started synthesizing Zx but, different from WT, its accumulation already saturated after approximately 30 min. In VDE-OE and Z1-V-OE, Zx formation was instead massively accelerated and saturated after approximately 30 min, reaching maximal levels similar to the WT (Figure 2B).

Concomitantly with Zx formation, we observed a corresponding decrease in Vx, which was clearly faster in VDE-OE and Z1-V-OE cells. In those lines, the decrease in Vx plateaued after approximately 30 min, and WT cells eventually reached a similar Vx content at the end of the 90 min treatment. In ZEP1-OE cells, the decrease in Vx is similar rate than in WT but plateaued at higher levels (Figure 2C). WT and ZEP1-OE cells accumulated a considerable amount of semi-epoxidated Ax upon illumination. Interestingly, Ax in ZEP1-OE cells reached higher levels than in WT, implying that despite strong XC activation in light, ZEP1 overexpression could counteract endogenous VDE activity and produce Ax from previously synthesized Zx. Instead, the VDE-OE and Z1-V-OE lines accumulated less Ax, suggesting that the activity of VDE is dominant over ZEP1 and did not allow the accumulation of this partially de-epoxidated xanthophyll species (Figure 2D).

Upon exposure to dark after the light treatment, the XC relaxed, starting with different initial xanthophyll compositions in the four lines due to the different kinetics of activation (Figure 2B-D, S6E). Zx was consumed at a constant and similar rate in WT and VDE-OE cells. Instead, in the two lines that overexpress ZEP1, Zx was reconverted faster (Figure 2B). The restoration of the Vx content showed a consistent trend, with a similar synthesis rate in WT and VDE-OE cells but faster rates in the ZEP1-OE and Z1-V-OE lines (Figure 2C). The Ax dynamics in the dark showed a more complex picture, indicative of this carotenoid being produced simultaneously from Zx and consumed to synthesize Vx. Upon switching from light to darkness, the WT, Z1-V-OE, and VDE-OE lines all initially showed elevated Ax levels, consistent with their increased Zx content. In general, Ax was reconverted more rapidly in lines with an increase in the content of ZEP1 (Figure 2D).

Taken together, these results show that increased accumulation of ZEP1 and VDE can improve XC dynamics. A relatively low increase in enzyme accumulation (2-3 times higher than in WT cells) had a large impact on xanthophyll dynamics, demonstrating that both enzymes are rate-limiting components of the xanthophyll cycle in *N. oceanica*.

### Alteration of xanthophyll cycle dynamics strongly affects NPQ kinetics

Like other heterokont algae, Zx accumulation has a strong impact on NPQ activation and relaxation in *Nannochloropsis* (Bna et al., 2017), thus, we characterized how changes in XC dynamics affected NPQ kinetics upon illumination with different light protocols. Upon exposure to saturating light, ZEP1 overexpression led to lower maximal NPQ levels (Figure 3A). This difference was lost using oversaturating actinic light (Figure 3B), indicating that excess illumination could outweigh ZEP1 overexpression by increasing activation of endogenous VDE. However, in both light conditions, NPQ relaxation was significantly faster in ZEP1-OE compared to WT. Instead, overexpression of VDE caused massively accelerated NPQ activation upon exposure to saturating actinic light. The NPQ capacity was already close to saturation after 30 s of illumination, while it took approximately 10 min for the WT to reach similar levels (Figure 3A). This result is consistent with the acceleration of Zx formation in VDE-OE cells. Expectedly, the effect was also present upon illumination with oversaturating light (Figure 3B). In both kinetics, the relaxation of NPQ in darkness was much slower in VDE-OE than in the WT, which is in accordance with the slow Zx reconversion rates in the VDE-OE lines observed before. In fact, even after a relatively short actinic light treatment of 10 min, the VDE-OE cells fully relaxed NPQ in 25 min, while WT relaxed almost immediately (Figure 3C).

**Figure 3.**
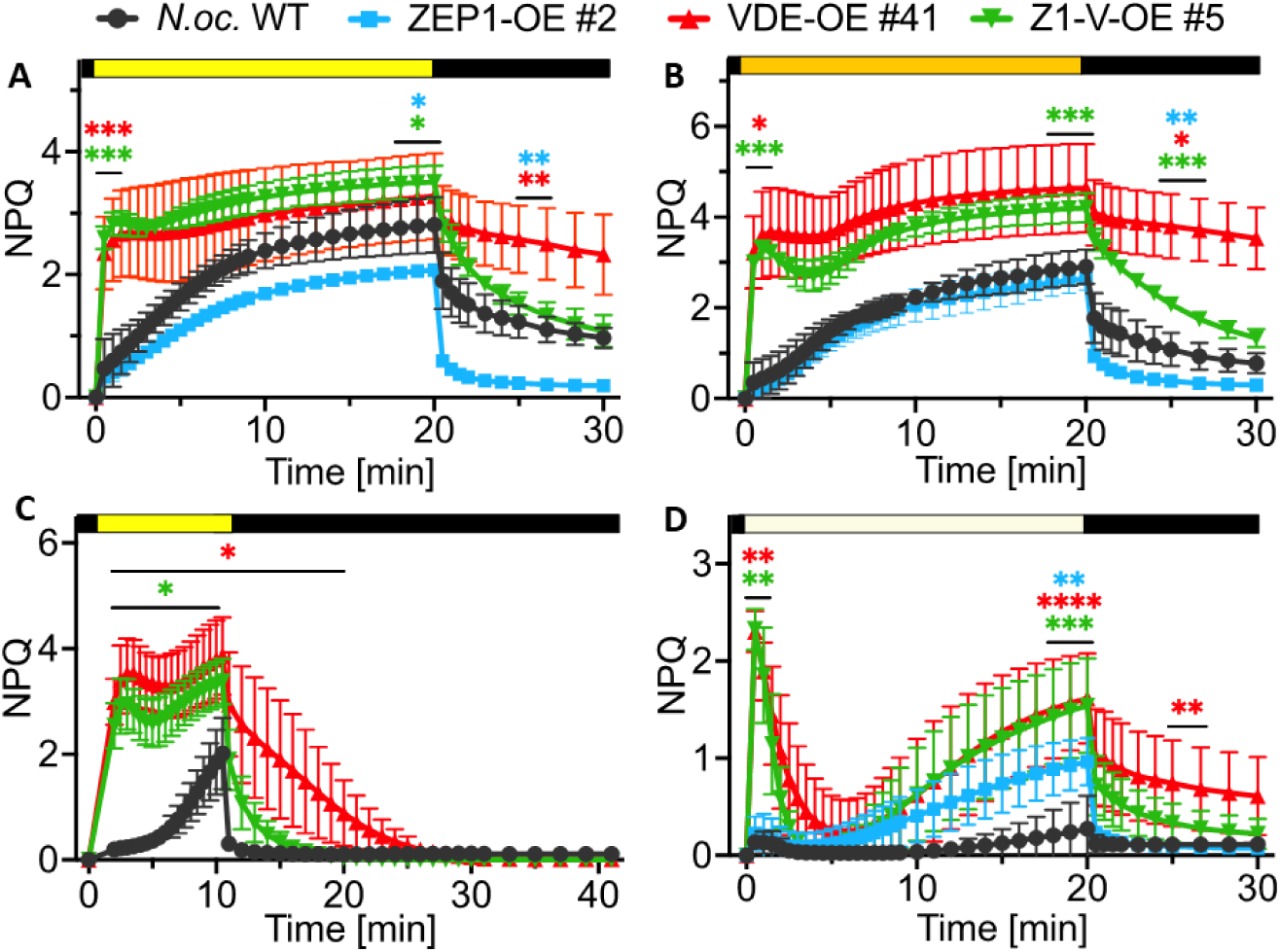
Non-photochemical quenching (NPQ) kinetics in ZEP1/ VDE overexpression lines under varied light treatments. N.oc. WT and Lines overexpressing ZEP1/ VDE were dark-adapted, subjected to different light treatments and NPQ kinetics were assessed. A) NPQ activation for 20 min under saturating Light (1000 µmol photons s^-1^ m^2^), followed by a dark phase of 10 min. B) NPQ activation for 20 min under over-saturating light (2000 µmol photons s^-1^ m^2^), followed by 10 min darkness. C) NPQ activation for 10 min at saturating light (1000 µmol photons s^-1^ m^2^), followed by 30 min darkness. D) NPQ activation for 20 min under non-saturating light (175 µmol photons s^-1^ m^2^), followed by 10 min darkness. Data is represented as mean ± SD from ≥ 3 independent biological replicates. For A), B) and D), significant differences from the WT were assessed with one-way Welch-ANOVA tests for the first 2 time points after switching on actinic Light, the last 3 time points in actinic light and 2 time points during the dark phase. For C), significance was assessed for each data point using two-tailed Student’s t-tests. *p < 0.05, **p < 0.01, ***p < 0.001, ****p < 0.0001.

Z1-V-OE cells that accumulate both ZEP1 and VDE remarkably showed a combined phenotype between the single overexpressing lines. In the light phase of all treatments, the effect of VDE overexpression was dominant, and the cells showed extremely fast activation of NPQ, as seen in VDE-OE cells. When switched to the dark, NPQ relaxation was markedly faster than in VDE-OE cells (Figure 3A-C), suggesting that ZEP1 accumulation was effective. These results are consistent with the accelerated kinetics of XC mentioned above in Z1-V-OE cells and show that the dynamics of the XC are seminal for NPQ activation in *N. oceanica*, with VDE/ ZEP activity having a major impact on controlling this regulatory response.

Upon exposition to non-saturating actinic light NPQ was massively activated in VDE-OE and Z1-V-OE cells. In 30 s of low illumination, NPQ reached values close to those observed with much stronger illumination (approx. NPQ = 2.5), while in WT cells NPQ was almost not activated (NPQ = 0.14). However, after initial activation we observed a strong drop in the level of NPQ, which can be explained by the activation of the Calvin-Benson cycle after a few minutes, acting as an electron sink and dissipating a part of the lumenal ΔpH, relaxing the NPQ (Figure 3D).

In the same condition, the initial activation of NPQ in ZEP1-OE was similar to the WT, while it increased over time to a higher steady state (Figure 3D). This can be explained by observing that dark-adapted ZEP1-OE cells contained less Zx than WT (Figure S6D) and thus the WT cells were already in a partially quenched state. Since NPQ quantifies the difference between dark-adapted and illuminated cells, the quenching induced by the dim light treatment is thus larger in ZEP1-OE than in WT because they start from a different initial state. In addition, in this case, the NPQ in ZEP1-OE recovered very quickly after the light was turned off (Figure 3D).

The observed NPQ phenotypes of the strains ZEP1-OE, VDE-OE, and Z1-V-OE were confirmed in multiple independent lines that were indistinguishable in all performed analyses (Figure S7). As the effect of ZEP1 overexpression on photosynthesis and NPQ was highly impactful, we revisited the *zep2* gene in *N. oceanica,* generating ZEP2 overexpression lines (ZEP2-OE) and characterizing their NPQ dynamics. Induction of NPQ under saturating light in ZEP2-OE was comparable to that of the WT, while relaxation of NPQ in darkness was accelerated, confirming its function as a ZEP (Figure S8). However, as the effect of overexpression seemed to be less strong, we decided to continue with the ZEP1-OE line only.

Changes in ZEP1 and VDE content also had effects on photosystem II (PSII) photosynthetic efficiency during illumination. Under saturating actinic light, there was no clear difference between the overexpression strains and the WT, as the photosynthetic capacity expectedly was saturated in all lines. However, recovery of photosynthetic efficiency in the darkness was faster in ZEP1-OE lines (Figure 4, S9A) and slower in the VDE-OE lines (Figure 4, S9B), suggesting that higher photoprotection caused a penalty in photosynthetic efficiency when light was turned off. The Z1-V-OE lines were very similar to the WT (Figure 4, S9C). These trends show that XC regulation has a notable impact on photochemical activity and is not limited to photoprotection alone.

**Figure 4.**
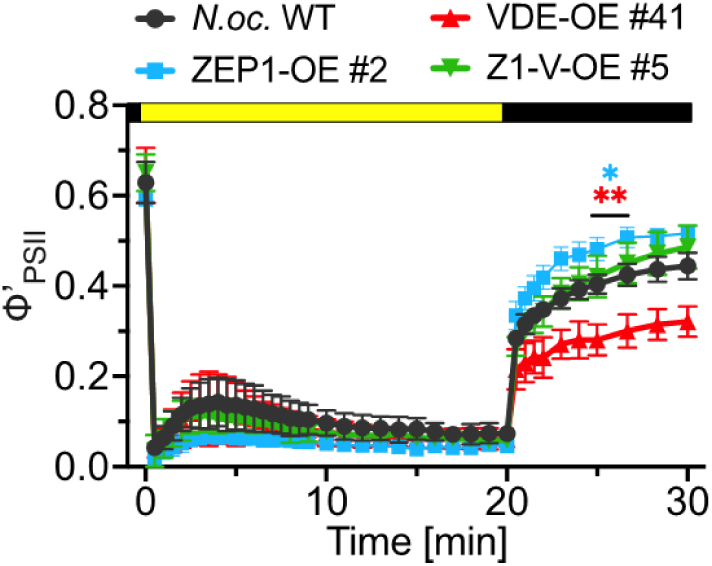
Photosynthetic efficiency (φ’_pSII_) of ZEP1 / VDE overexpression strains. *N. oc.* WT and the strains generated in this study were dark-adapted and exposed to 20 min of high light (1000 µmol photons s^-1^ m^2^), followed by 10 min in darkness. Significant differences to the WT were assessed for the two points during the dark phase using one-way ANOVA tests, *p < 0.05, **p < 0.01.

### The accumulation of ZEP1 / VDE strongly impacts the response to light fluctuations

Since ZEP1 / VDE overexpression showed a substantial impact on XC dynamics and NPQ activation, we next sought to analyze the impact of faster light changes, which *N. oceanica* could encounter in nature due to rapidly changing weather conditions and/ or turbulences in the water column.

For this, the strains were exposed to alternating saturating light and dark phases with different kinetics while monitoring chlorophyll fluorescence and NPQ. First, we subjected the cells to cycles of 8 min in light and 10 min in darkness (Figure 5A). As described in the literature, we observed that the activation of NPQ in WT and ZEP1-OE cells increased with each cycle (Short et al., 2022). Because the dark phases were not long enough to fully epoxidate Zx, cells continued to accumulate it, leading to an increase in NPQ over time. In VDE-OE and Z1-V-OE, this effect was smaller, as the Zx generation was so fast that NPQ was already maximally activated in the first light cycle.

**Figure 5.**
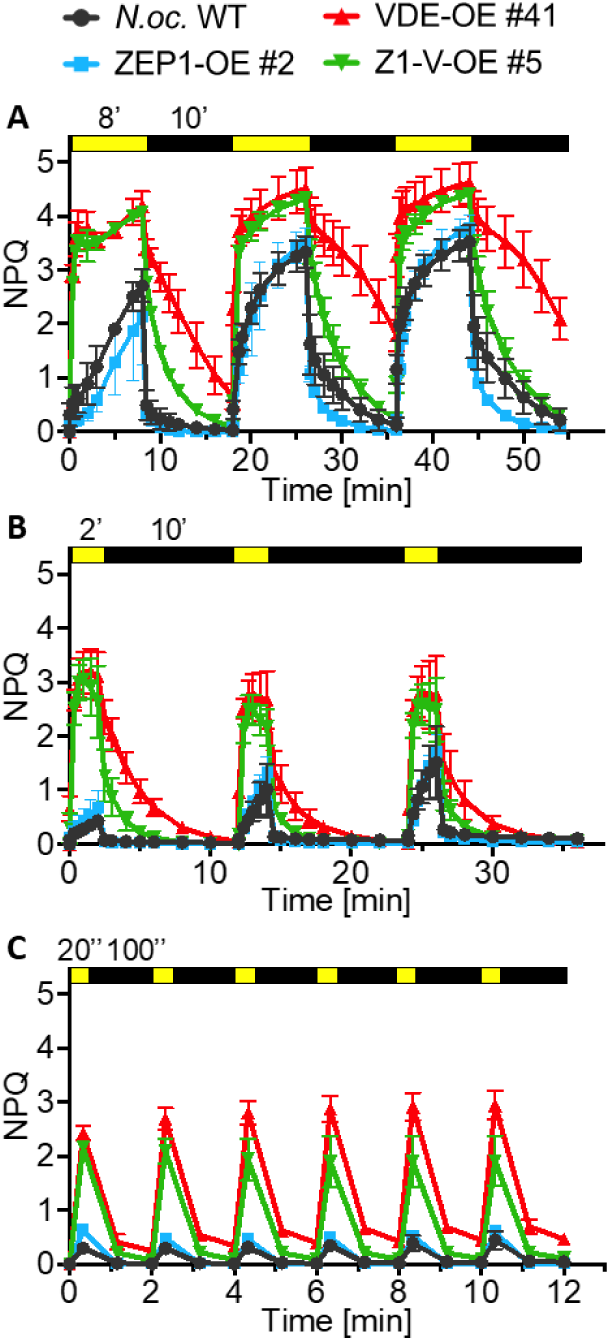
Impact of ZEP1 / VDE overexpression on NPQ kinetics under fluctuating light. Dark-adapted (30 min) cells of the three transgenic lines and *N.oc.* WT were exposed to different light/ dark sequences and NPQ kinetics were monitored. A) Fluctuation of 8 min light and 10 min darkness. B) Fluctuation of 2 min light and 10 min darkness. C) Fluctuation of 20 s light and 100 s darkness. NPQ was measured right before actinic light was turned on and off and in the middle of the dark phase. Time intervals are indicated above each graph, ’minutes, "seconds. In all figures data is represented as mean ± SD from ≥ 3 independent biological replicates.

However, VDE-OE showed only partial recovery in the dark. This difference increased with every cycle, suggesting that the high activity of VDE caused the progressive accumulation of Zx. This was not the case for Z1-V-OE, which almost fully relaxed NPQ after each cycle, suggesting that increased ZEP1 was able to balance the higher activity of VDE during the dark phases.

To analyze the effect of shorter light phases, we reduced the time of exposure to actinic light to 2 min, maintaining the same dark duration (Figure 5B). Under this regime, the illumination time was not sufficient for the ZEP1-OE and WT lines to fully activate NPQ. The two lines were also indistinguishable in their dark relaxation phase, as the endogenous ZEP1 was presumably sufficient to reconvert Zx and relax the XC and NPQ in the WT after the shorter light phase. The lines with higher VDE accumulation were instead both able to fully activate NPQ, even upon a shorter illumination time. In the dark, NPQ relaxed faster in the Z1-V-OE line, but 8 min in the dark were also sufficient for complete relaxation of VDE-OE (Figure 5B).

We also tested even faster dynamics with 20 s of light and 100 s of darkness (Figure 5C). After a few cycles, the maximum NPQ levels after light exposure stabilized in all lines. Fascinatingly, even in these very short light pulses, Z1-V-OE and VDE-OE cells could activate a strong NPQ response, whereas NPQ activation in WT and ZEP1-OE was close to zero. This demonstrates that with fast light fluctuations, stronger accumulation of VDE makes the difference in activating a significant photoprotective response in *N. oceanica*.

### A balanced xanthophyll cycle contributes to ROS scavenging

The role of Zx in photoprotection is not restricted to NPQ, but it also increases the cell’s ability of scavenging ROS, a finely regulated and strain-specific feature (Sáenz et al., 1997; Vonshak et al., 2020; Ben-Sheleg and Vonshak, 2022). XC manipulation could affect this response, and for this reason we assessed the sensitivity of the four strains to ROS by exposing them to ROS producing agents and following their growth. The algae cultures were grown at 100 µmol photons m-2 s-1 since we saw that this light intensity was sufficient to induce differences in Zx accumulation in transgenic lines (Fig. 2A), but it was not saturating (Vonshak et al., 2020). First, we added methylviologen (MV) to induce the formation of superoxide anions (O2•-). Since the additional O2•-was externally generated (i.e., not in the photosynthetic apparatus) and is only quenched chemically, any potential effect should be associated with the ROS scavenging ability and not with the modulated energy quenching activity in the different overexpression lines. Upon MV treatment, the growth of all strains was reduced, confirming the toxicity of the generated O2•-(Figure 6A-B). As MV is constantly reduced in the chloroplast, it was expected to be maintained throughout the duration of the experiment. Interestingly, after 5-6 days, we were able to observe that the growth of the Z1-V-OE strain was significantly higher than that of WT cells (Figure 6B), suggesting the increased ability of Z1-V-OE to withstand ROS. The ZEP1-OE line, on the other hand, showed reduced growth, reflecting increased ROS sensitivity. The VDE-OE line was less affected and grew similarly to the WT. Overall, these data suggest that reducing Zx accumulation could decrease resistance to O2•-, while increasing XC turnover rates might be beneficial.

**Figure 6.**
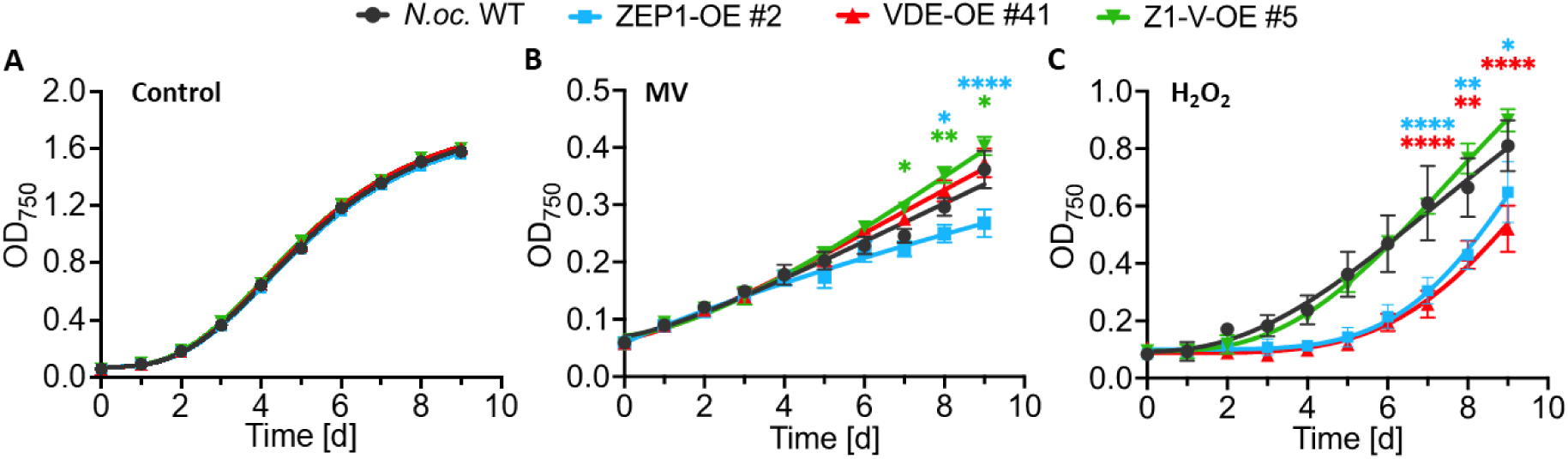
Sensitivity of *N. oceanica* lines to chemically induced oxidative stress. *N. oc.* WT and lines of ZEP1-OE, VDE-OE and Z1-V-OE were cultivated under constant illumination (100 µM photons s^-1^ m^2^), and in 1% CO_2_. **A)** Controls were grown without additional ROS producers. **B)** Formation of superoxide was induced by addition of 50 µM methyl viologen (MV). **C)** Resistance to hydroxen peroxide was determined by adding 0.5 mM H_2_O_2_. Sensitivity to ROS was assessed by observing growth for 9 days measuring OD_750_. Significant differences from *N.oc.* WT were determined with one-way Welch-ANOVA tests on the last three days for each experiment, *p < 0.05, **p < 0.01, ****p < 0.0001. Data is represented as means ± SEM from s 4 biological replicates. Trendlines represent fittings to a sigmoidal growth model.

In chloroplasts, O2•- is also partially disposed of by superoxide dismutase, which generates hydrogen peroxide. To dissect whether the observed phenotype upon MV addition was caused by actual accumulation of O2•- or due to secondary H2O2 production, we directly tested the effect of H2O2 on algal growth. Unlike MV, H2O2 is consumed over time, which means that most stress was limited to the first few days after treatment. ZEP1-OE cells again exhibited reduced growth, whereas the Z1-V-OE line was similar to the WT. Surprisingly, VDE-OE cell growth was delayed by H2O2 addition (Figure 6C), suggesting that an increase in Zx is not necessarily beneficial if XC is imbalanced. This also reveals that the growth phenotype observed under the addition of MV was indeed mainly caused by stress and most likely not related to secondary H2O2.

Overall, these results underline that the XC impacts tolerance to different ROS and that disequilibrating this delicately balanced mechanism can have detrimental effects.

### Impact of XC modulation on growth under variable light conditions

The effects of XC modulation on NPQ kinetics and ROS clearance suggest that the strains generated in this work might also show a growth phenotype under challenging light conditions. Therefore, lines overexpressing ZEP1/ VDE were tested for their growth on plates exposed to various light regimes. Under low light conditions (LL, 25 µmol photons m-2 s-1), all four lines showed the same growth, which could be expected considering that the XC should not be activated under low light (Figure 7A-B). When cells were exposed to higher, saturating, illumination (HL, 100 µmol photons m-2 s-1 for one week followed by one week in 2000 µmol photons m-2 s-1), ZEP1-OE cells exhibited slightly defected growth (Figure 7A-B). The VDE-OE line grew denser than the WT, highlighting that the increased Zx accumulation caused to overcome the limit of light stress tolerance. Z1-V-OE cells exhibited a growth similar to the WT, reflecting that the accumulation of ZEP1 counteracted VDE, leading to a similar XC balance. To assess how XC modulation affected growth under dynamic light conditions, we exposed the strains to fluctuating light (FL 1, 500 µmol photons m-2 s-1 with 1:4 min light: dark cycles), which can be particularly stressful for *Nannochloropsis* (Bellan et al., 2020). Under this regime, VDE-OE and Z1-V-OE cells showed growth close to WT cells, while ZEP1-OE cells were largely affected, showing a major growth reduction (Figure 7A-B).

**Figure 7.**
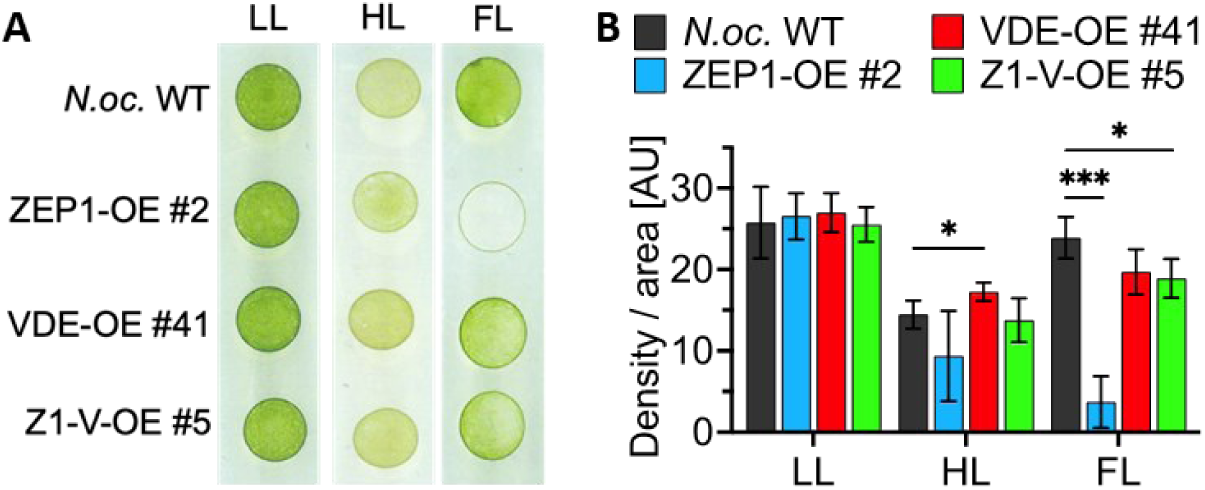
On-plate characterization of N. oceanica lines growing under different light regimes. 10^6^ cells of N.oc. WT and random lines of ZEP1-OE, VDE-OE and Z1-V-OE were plated on F/2 supplemented with 10 mM sodium bicarbonate and grown under different light treatments for 2 weeks (LL, HL) or 4 weeks (FL). A) Growth phenotype of strains treated with LL, low light (16:8 h light:dark photoperiod in 25 µmol photons m^2^ s^1^); HL, high light (16:8 light:dark regime under 100 µmol photons m 2 s’^1^ for one week, followed by 2000 µmol photons m^-2^ s’^1^ for 1 week) and FL, fluctuating light (fluctuations of 1 min light (500 µmol photons m^2^ s^-1^) and 4 min darkness) B) Quantification of spot density per area from plates shown in A). Quantification was performed with Fiji/ ImageJ. Data is represented as mean ± SD from 4 biological replicates. Significance was assessed with pairwise one-tailed Student’s t-tests for each individual light treatment, *p < 0.05, ***p < 0.001. AU, arbitrary units.

## DISCUSSION

### The VDE and ZEP1 activities determine the dynamics of XC in Nannochloropsis

The seawater alga *Nannochloropsis* exhibits a very high violaxanthin content (up to 3.9 mg / g of biomass), probably the highest among photosynthetic species investigated so far (Lubián et al., 2000; Basso et al., 2014; Park et al., 2021). Similarly to other secondary endosymbiotic algae such as diatoms, *Nannochloropsis* strongly relies on the XC for the regulation of photosynthesis (Bína et al., 2017; Lacour et al., 2020; Zainal Abidin et al., 2021).

To assess the impact of XC on photoprotection and photosynthesis regulation in *Nannochloropsis oceanica,* in this work we generated strains with altered accumulation of VDE and ZEP1. The modified strains showed a strong alteration in the XC dynamics (Figure 2). With higher VDE accumulation, the speed of de-epoxidation of Vx was massively increased (Figure 2C), revealing that the de-epoxidation rate is directly regulated by the enzyme activity and modulation of the enzyme content is the main determinant for the zeaxanthin accumulation rate.

This is different from plants, where the XC rate is principally determined by the availability of ascorbic acid (Neubauer and Yamamoto, 1994; Müller-Moulé et al., 2002), and the release of the xanthophylls from the antenna complexes (Jahns et al., 2009). There is evidence that VDE activity can also be regulated by protein levels in plants (Bugos et al., 1999; Zhu et al., 2017), however, overexpression studies in tobacco and *Arabidopsis* revealed that even strongly enhanced VDE accumulation yielded only minorly increased de-epoxidation rates (Hieber et al., 2002; Deng et al., 2003). Thus, while in plants the XC is mainly regulated by the availability of its substrates violaxanthin and/ or Asc, in *N. oceanica* the main limiting factor is the enzyme activity, showing how during evolution different organisms can exploit and optimize the same tools differently.

Interestingly, VDE overexpression drastically impacted the kinetics of Zx synthesis, but it did not alter the maximal levels of Vx de-epoxidation, which were similar in WT and VDE-OE (Figure 2C), consistent with observations in diatoms (Manfellotto et al., 2020; Küster et al., 2023) and plants (Chen and Gallie, 2012). In photosynthetic eukaryotes, xanthophylls are mostly bound to antenna proteins, where they can be exchanged and de-epoxidated (Morosinotto et al., 2002). However, a fraction of Vx is stably bound to antenna binding sites that are unavailable for the VDE activity (Wehner et al., 2006). In *Nannochloropsis* WT cells, approximately 65% of Vx is available for de-epoxidation, while the rest is not converted into Zx even if exposed to extreme illuminations (Perin et al., 2023). This behavior is also congruent with observations in lines of *Arabidopsis* with an increased xanthophyll pool (Johnson et al., 2008), suggesting that maximum Vx de-epoxidation is not determined by the Vx / VDE ratio, but by physical constraints in the binding sites within the photosynthetic apparatus.

The increased accumulation of ZEP1 also showed a clear impact on XC dynamics, which is particularly evident in the dark after light treatment (Figure 2). Cells overexpressing ZEP1 showed faster recovery after light was turned off, with quicker re-synthesis of Vx, independent of the genetic background (WT or VDE-OE) or the illumination conditions tested (Figure 2C). This clearly shows that, as with VDE, the ZEP1 content is a major factor in the control of the XC kinetics in *N. oceanica.* The impact of ZEP1 overexpression was also measurable during exposure to saturating light. Here, ZEP1-OE cells showed lower levels of Zx than the parental line (Figure 2), demonstrating that while the activity of VDE is dominant and drives the massive formation of Zx when activated by low lumenal pH, ZEP is also active under illumination and affects the balance of the XC. ZEP1 overexpression decreased Zx accumulation in the light with a consequent reduction in photoprotection capacity (Figure 3A) and tolerance to light stress (Figure 7).

When VDE and ZEP1 were overexpressed together, the cells showed faster Zx synthesis and reconversion in the dark (Figure 2B). This suggests that the activities of the two enzymes are rather independent and that, apart from the shared substrate pool, there is no significant reciprocal influence. This is consistent with the localization of the enzymes on the stromal (ZEP1) and the lumenal (VDE) side, while the substrates are found in the thylakoid membrane. Western blot analysis also suggests that while VDE is a soluble protein that associates with the membrane only upon activation (Arnoux 2009), ZEP1 in *Nannochloropsis* is likely to stably attach to the membrane, a property that is not shared by all ZEP isoforms in plants (Schwarz et al., 2015; Bethmann et al., 2019). VDE and ZEP1 also share a substrate, antheraxanthin (Ax), which acts as an intermediate of the conversions between Vx and Zx. Ax itself is also expected to contribute to photoprotection (Goss et al., 1998; Short et al., 2023), even though this hypothesis is debated (Bína et al., 2017). In the two lines with overexpressed VDE, the accumulation of Ax was reduced during light exposure, while in the ZEP1-OE lines its accumulation was slightly promoted (Figure 2D), suggesting that Ax is mostly generated from Zx by ZEP1 and removed by VDE.

It is worth underlying that the impact of the VDE/ ZEP1 accumulation levels on XC dynamics is observed not only upon saturating illumination but also in cells grown under non-stressful dim light conditions (e.g. 100 µmol photons m-2 s-1). These cells showed in fact a threefold increase in Zx accumulation in the VDE-OE line, while it was one third lower in ZEP1-OE (Figure 2A), suggesting that XC is already involved in the regulation of photosynthesis under relatively mild light in *N. oceanica*.

A comparative study in six algal species proposed a range of the light threshold for XC activation from 80 – 170 µmol photons m-2 s-1 (Dimier et al., 2009), classifying *N. oceanica* among those that respond quickly. These observations support the idea that the XC in *N. oceanica* should not simply be considered as a mechanism to respond to excess illumination but rather a central player in the modulation of photosynthesis efficiency and photoprotection under all conditions.

### The xanthophyll cycle controls NPQ activation in Nannochloropsis

The alterations of the XC dynamics in the analyzed lines have a major impact on the NPQ kinetics. The VDE-OE strains under various light conditions showed an almost immediate NPQ response, whereas the ZEP1-OE lines had reduced steady-state NPQ levels under saturating actinic light and were faster at relaxing. Simultaneous overexpression of VDE and ZEP1 led to an additive effect with an overall acceleration of XC in both directions (Figure 3, S7).

NPQ in *N. oceanica* is known to be strongly dependent on the XC (Bína et al., 2017). This work also allows quantitative comparison of NPQ levels with the de-epoxidation state 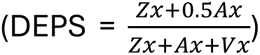 (Figure 8), showing that there is a strong correlation between DEPS and NPQ levels. However, this correlation is only valid until NPQ saturates around values of 2-3 and DEPS of 0.1-0.2, depending on the ZEP1/ VDE content. For higher DEPS values, increasing de-epoxidation no longer corresponds to higher NPQ levels (Figure 8).

**Figure 8.**
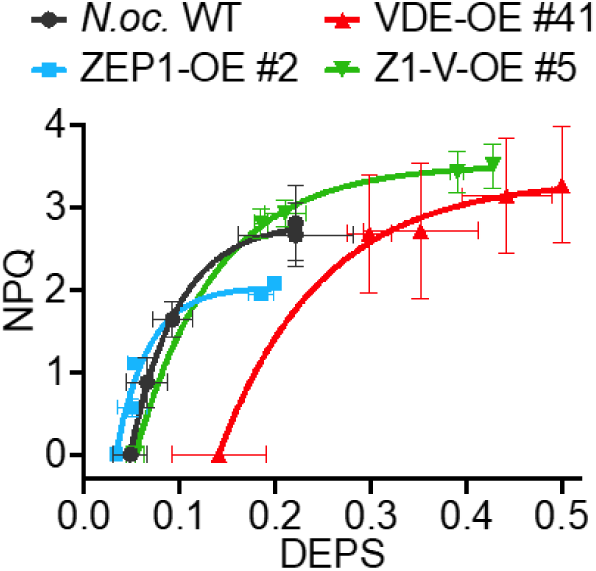
Correlation of de-epoxidated xanthophyll species (DEPS) and NPQ. Levels of the de-epoxidated xanthophyll states 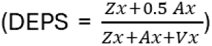 in N.oc. WT and randomly selected lines of Zx+Ax+Vx ZEP1-OE, VDE-OE and Z1-V-OE were plotted against NPQ levels after exposition to light periods of 0, 2, 5, 15 and 20 min at 1000 µmol photons m ^-2^ s^-1^ (Values extracted from Figures 2B-D and 3A). DEPS levels at 20 min were extrapolated from fitted curves in Figure 2B-D and plotted against NPQ levels after 20 min in light. Data points are represented as mean ± SD for both DEPS and NPQ. Curves show fittings to exponential curves.

In *Nannochloropsis*, similarly to other eukaryotic algae, NPQ activation also requires the presence of specialized antenna proteins of the LHCX family (Li et al., 2000; Buck et al., 2019; Park et al., 2019). One possible mechanistic explanation for the observed biphasic correlation between NPQ and DEPS is that LHCX must bind Zx to induce quenching, but there is a limit at which the available binding sites in LHCX are saturated, and thus, the additional synthesis does not induce any increase in quenching.

When comparing the results obtained with other photosynthetic eukaryotes, a fully linear correlation between DEPS and NPQ has been shown in some diatoms (Dimier et al., 2009; Stamenković et al., 2014; Giossi et al., 2024). In the green alga *Chlorella vulgaris*, which relies strictly on XC for photoprotection too, the DEPS-NPQ relation shows the same asymptotic behavior that we observed for *N. oceanica* (Quaas et al., 2015; Girolomoni et al., 2020), indicating that the XC-NPQ relation could be the result of convergent adaptation rather than phylogenetic connection.

This idea is consistent with comparative analyses of different algal ecotypes that show how regulation of the XC and quenching efficiencies is generally a feature associated with ecological adaptation rather than with evolutionary trends (Barnett et al. 2015; Quaas et al. 2015; Lacour et al. 2020). *Nannochloropsis* species were found in diverse environments around the world (Suda et al. 2002; Sandnes et al. 2005; Fawley et al. 2015; Karlov et al. 2017), probably needing different optimization of mechanisms to regulate photosynthesis. Our work revealed that NPQ activity in this species can be efficiently regulated by modulating the VDE and ZEP1 content, affecting the light threshold for NPQ induction and maximum NPQ levels. The control of their expression could thus be a strategy for tuning the XC and, consequently, adjusting photosynthesis regulation to the specific environmental niche.

### The role of ZEP in the xanthophyll cycle is crucial for balancing photoprotection and light use efficiency

The XC has a major impact on NPQ kinetics, but its biological role is wider. The accumulation of Zx contributes to the scavenging of toxic ROS (Havaux et al., 2007; Müller et al., 2011), determining the photoprotective potential. Under externally generated superoxide anions (O2•-) and hydroxide peroxide (H2O2), the ZEP1-VDE double overexpressor showed partially improved resistance, while the single overexpression of ZEP1 or VDE resulted in a general decrease in ROS tolerance (Figure 6). Both ROS mainly react with Zx to form various Zx epoxides that can be recycled in part by the XC (Nishino et al., 2017). Acceleration of the XC might improve clearance of these epoxide species and therefore regeneration of protective Zx, rescuing growth defects. However, chemical induction of ROS could also induce oxidative stress in other compartments or trigger potential ROS signaling pathways (Ben-Sheleg and Vonshak, 2022), requiring a deeper investigation of the redox regulation in *Nannochloropsis*.

The impact of XC modulation on the tolerance of NPQ and ROS consequently affected growth of the algal strains under various light regimes. While the XC does not majorly influence growth under low light, its importance became evident under more stressful high light and fluctuating light conditions, in particular showing that strains overexpressing ZEP1 showed higher photosensitivity, associated with a reduced capacity to accumulate Zx and activate NPQ (Figure 7).

These observations highlight the fact that, despite VDE playing an outstanding role in XC activation, ZEP activity also has a major, and often underestimated, impact on regulation and photoprotection. The reduction in the maximum achievable levels of Zx and NPQ in ZEP1-OE under saturating light suggests that ZEP1 controls the steady-state equilibrium of the XC, making cells more susceptible to stressful light conditions. This is in line with studies in tobacco and *Arabidopsis*, which reveal that ZEP content regulates maximum de-epoxidation levels and can indeed confer increased light sensitivity upon overexpression (Wang et al., 2008; Küster et al., 2023).

Consistent with this idea, recent studies presented regulatory mechanisms of ZEP activity in plants, which was proposed to be activated in light through redox regulation via the chloroplast NTRC and thioredoxin system (Naranjo et al., 2016; Da et al., 2018). ZEP activity was also shown to be inhibited by H2O2 (Holzmann et al., 2022) or by inducing ZEP degradation upon prolonged light stress (Bethmann et al., 2019), as strategies to maintain high Zx levels under those conditions.

The fact that ZEP activity increased photosensitivity is not necessarily negative. In fact, in plants and algae that excessive accumulation of zeaxanthin was shown to reduce photosynthetic efficiency (Kalituho et al., 2006; Liu et al., 2023). Thus, faster recovery from Zx accumulation could increase light-use efficiency under limiting light and even led to improved growth in plants and *Nannochloropsis gaditana* (Park et al., 2008; Perin et al., 2023).

These results demonstrate that accelerating Zx recovery is a valuable strategy to improve photosynthetic efficiency, but requires delicate fine tuning, as excessively reduced levels of Zx can diminish the cell’s ability to withstand light stress. Furthermore, they show that modulation of the ZEP activity can be a simple and useful strategy for adapting to rapidly changing light conditions in the dynamic habitats of *Nannochloropsis* (Owens et al., 1987; Ashour et al., 2019; Canini et al., 2024).

## Acknowledgments

Tim M. and TM acknowledge funding from European Union H2020 Marie Skłodowska-Curie No. 955520 Digitalgae. Authors thank Sarah D’Adamo (Wageningen University, The Netherlands) for *N. oceanica* tdTomato strain.

